# *tidysdm*: leveraging the flexibility of *tidymodels* for Species Distribution Modelling in R

**DOI:** 10.1101/2023.07.24.550358

**Authors:** Michela Leonardi, Margherita Colucci, Andrea Vittorio Pozzi, Eleanor M. L. Scerri, Andrea Manica

## Abstract

In species distribution modelling (SDM), it is common practice to explore multiple machine-learning algorithms and combine their results into ensembles. This is no easy task in R: different algorithms were developed independently, with inconsistent syntax and data structures. Specialised SDM packages integrate multiple algorithms by creating a complex interface between the user (providing a unified input and receiving a unified output), and the back-end code (that tackles the specific needs depending on the algorithm). This requires a lot of work to create and maintain the right interface, and it prevents an easy integration of other methods that may become available.

Here we present *tidysdm*, an R package that solves this problem by taking advantage of the *tidymodels* universe. Being part of the *tidyverse*, (i) it has standardised grammar and data structures providing a coherent interface for modelling, (ii) includes packages designed for fitting, tuning, and validating various models, and (iii) allows easy integration of new algorithms and methods.

*tidysdm* allows easy, flexible and quick species distribution modelling by supporting standard algorithms, including additional SDM-oriented functions, and giving the opportunity of using any algorithm or procedure to fit, tune and validate a large number of different models. Additionally, it provides further functions to easily fit models based on paleo/time-scattered data.

The package includes two vignettes detailing standard procedures for present-day and time-scattered data. These vignettes also showcase the integration with *pastclim* (Leonardi *et al*. 2023) to allow easier access to palaeoclimatic data series, if needed, but users can bring in their own climatic data in standard formats.

## INTRODUCTION

Species Distribution Modelling (SDM, also known as Ecological Niche Modelling, ENM; Habitat Suitability Models, HSM; and a few other acronyms (Guisan *et al*. 2017)) is a class of biological methods that uses the occurrences of a species and the associated environmental variables to predict how suitable a geographic area is for the species itself (Elith and Leathwick 2009). SDMs have been used for a variety of scopes, *e*.*g*. to predict the distribution of invasive species in a new range (Elith 2017), to assess the habitat suitability for a given organism under future climatic scenarios (Franklin 2023), to reconstruct the geographic range of species in the past based on palaeontological remains (e.g. Leonardi *et al*. 2018) or present-day surveys (e.g. Miller *et al*. 2021b) and to detect changes in the realised niche of a species (e.g. Leonardi *et al*. 2022).

In computational biology, the field of SDM has been among the first ones to take advantage of machine learning, and it is now standard practice to explore multiple algorithms and combine their results into ensembles (Araújo and New, 2007). The R language (R core Team, 2023) is the most commonly adopted framework to fit SDMs, but it comes with the challenge that individual machine learning algorithms have been developed independently, with each package providing its own idiosyncratic way to format data and code the analysis. These differences make it difficult to repeat the same analysis with multiple algorithms and to create ensembles of their predictions. As a consequence, a large number of specialised SDM packages have been created (reviewed in Sillero *et al*., 2023), each attempting to provide a simple and unified interface to multiple algorithms. But this convenience is not without costs. The maintainers have to keep up with changes in a number of packages, which is resource intensive. Arguably more problematic is the fact that the package of choice limits the algorithms that are available while also constraining other approaches used in modelling (including the strategy used to create ensembles). Furthermore, innovations in the field of machine learning take longer to be adopted, as they need first to be implemented into a generic package, and then be ported into each of the specialised SDM ones.

The issues described above are not peculiar to SDMs, but to any field that leverages machine learning algorithms. This is why the *tidymodels* universe of packages (Kuhn and Wickham 2023) has emerged in R as a field-agnostic solution to these challenges, providing a standardised interface for modelling in R (*sklearn* provides a similar set of solutions in *python*). *tidymodels* includes packages specifically designed to fit, tune and validate a large number of different models. Its strength is given by the fact that it is embedded into the *tidyverse*, which philosophy is to have standardised grammar and data structures. In *tidymodels* every algorithm (no matter the library in which it is published) is provided with wrappers that accept the same input formats, use the same syntax, and produce the same output formats. In brief, it provides a coherent interface to modelling, while at the same time allowing the integration of any new algorithm, method or procedure that may be made available by the wider community. Packages such as *parnsip* (Kuhn and Vaughan 2023) and *yardstick* (Kuhn *et al*. 2023) provide the infrastructure to add respectively new models and metrics that seamlessly integrate within the broader *tidymodels* universe.

Here, we introduce *tidysdm*, a package that facilitates fitting SDMs with *tidymodels. tidysdm* integrates additional methods to several functions from the *tidymodels* universe to handle spatial data. It also implements algorithms and metrics favoured by SDMs practitioners but that are not available from other *tidymodels* packages. By building on the infrastructure of *tidymodels, tidysdm* takes advantage of a very large community of developers, rather than creating a complete solution from the ground up. Furthermore, as it follows all the standards *of tidymodels*, objects created within *tidysdm* can be fed to functions from a number of other packages, allowing, for example, for sophisticated stack ensembling out of the box. Any further expansion of functionality within the *tidymodels* universe will thus be immediately usable by SDM practitioners, without the need to wait for a field-specific implementation.

*tidysdm* also contains a number of functions to easily work with time-scattered data, a task that, with most other SDM packages, is not always possible or requires extensive tweaking. Furthermore, access and manipulation of (palaeo)climatic data and reconstructions are facilitated by integration with *pastclim* (Leonardi *et al*. 2023), a package designed to manipulate time series of palaeoclimate reconstructions. Recent changes in *pastclim* also allow it to easily download and manipulate present-day observations and future reconstructions, thus making it a helpful tool for any SDM application (but there is no requirement to use *pastclim;* the user can bring in any environmental data through their preferred pipeline, as *tidysdm* handles rasters with the widely used *terra* package (Hijmans 2023)).

We illustrate the functionality of *tidysdm* through two examples, one based on present-day occurrences of an endemic Iberian lizard, *Lacerta schreiberi*, to project its range in response to future climate change, and another reconstructing the range of wild horses through the Late Pleistocene and Early Holocene based on radiocarbon dates.

## EXAMPLE 1: SDM PIPELINE WITH PRESENT-DAY OCCURRENCES

The most common use of SDMs is to predict future distribution under global change from a set of presences in the present day. We provide an example workflow of such an analysis (https://evolecolgroup.github.io/tidysdm/articles/a0_tidysdm_overview.html), using presences for the Iberian emerald lizard, *Lacerta schreiberi*, an endemic species of the Iberian Peninsula (Rödder and Schulte 2010). The presences are provided as a dataset in *tidysdm* (they are also available as a GBIF Occurrence Download (6 July 2023) https://doi.org/10.15468/dl.srq3b3).

In *tidysdm*, presences are represented as spatial point objects using the library *sf* (Pebesma 2018; Pebesma and Bivand 2023). All functions in *tidysdm* can work with *sf* objects, and the package extends a number of methods from the *tidymodels* universe to handle them. Environmental raster data, on the other hand, are manipulated with *terra*. In this case, any further information specific to presences and absences/pseudo-absences/background is stored as additional columns in an *sf* object. *sf* and *terra* are arguably the most adopted spatial packages to represent point and raster data respectively.

An important challenge in most SDM analyses is that only presence data are available, and the choice of suitable pseudo-absences and the definition of the area of interest are key to informative modelling. These pre-modelling steps are outside the remit of *tidymodels*, but *tidysdm* does provide a number of functions to facilitate these tasks. We also note that there are many SDM packages (reviewed in Sillero *et al*. 2023) already available that can be used to complement the functions provided here. There is the caveat that not all of them are compatible with *sf* objects, but it is easy to convert an *sf* object to a plain data frame if necessary.

*terra* can be used to read environmental data in a number of formats. In the vignette, we use data from WorldClim (Fick and Hijmans 2017), a set of interpolated variables based on observations. There are a number of packages that help download WorldClim variables; in this example, we use *pastclim*, a library that is mostly designed to work on palaeoclimate time series, but also has a number of handy functions to download present-day data and future projections. *pastclim* returns standard SpatRaster objects from *terra*, thus allowing the use of standard *terra* functions to manipulate (crop, mask, project, etc.) the resulting raster. We use such functions in the example to focus our analysis on the Iberian peninsula.

A common challenge in SDMs is that the presences are not sampled in a structured manner over the whole region, but are the collation of haphazard efforts, leading to spatial sampling biases. A common solution to this problem is to thin the data (Steen *et al*. 2021), as done by the widely used package *spThin* (Aiello-Lammens *et al*. 2015). *tidysdm* contains functions to thin the dataset by distance (removing close-by points), as well as based on the cells of the raster (so that we only retain one presence per cell). By combining the functions *thin_by_dist() and thin_by cell()*, it is possible to prune datasets to remove (or at least minimise) spatial sampling biases (there are also analogous functions that work with time series of presences, *thin_by_dist_time(), thin_by_cell_time()*, see Example 2).

True absences are rarely available for analysis, as they require structured survey techniques with balanced sampling effort, whilst most datasets used for SDM are based on collations of opportunistic data. For this reason, a wide literature has arisen on how to sample pseudo-absences/background points (*e*.*g*. Barbet-Massin *et al*. 2012). *tidysdm* provides a number of commonly adopted techniques with the function *sample_pseudoabs():*

⍰ *random:* where the pseudo-absences/background are randomly sampled from the region covered by the raster (i.e. not NAs)
⍰ *dist_min:* pseudo-absences/background are randomly sampled from the region excluding a buffer of *dist_min* from presences (50 km in the example, shown in Figure 1a together with the thinned presences).
⍰ *dist_max:* pseudo-absences/background randomly sampled from the unioned buffers of *dist_max* from presences. Using the union of buffers means that areas that are in multiple buffers are not oversampled. This is also referred to as *“*thickening*”*.
⍰ *dist_disc:* pseudo-absences/background randomly sampled from the unioned discs around presences with the two values of *dist_disc* defining the minimum and maximum distance from presences.

**Figure 1:**
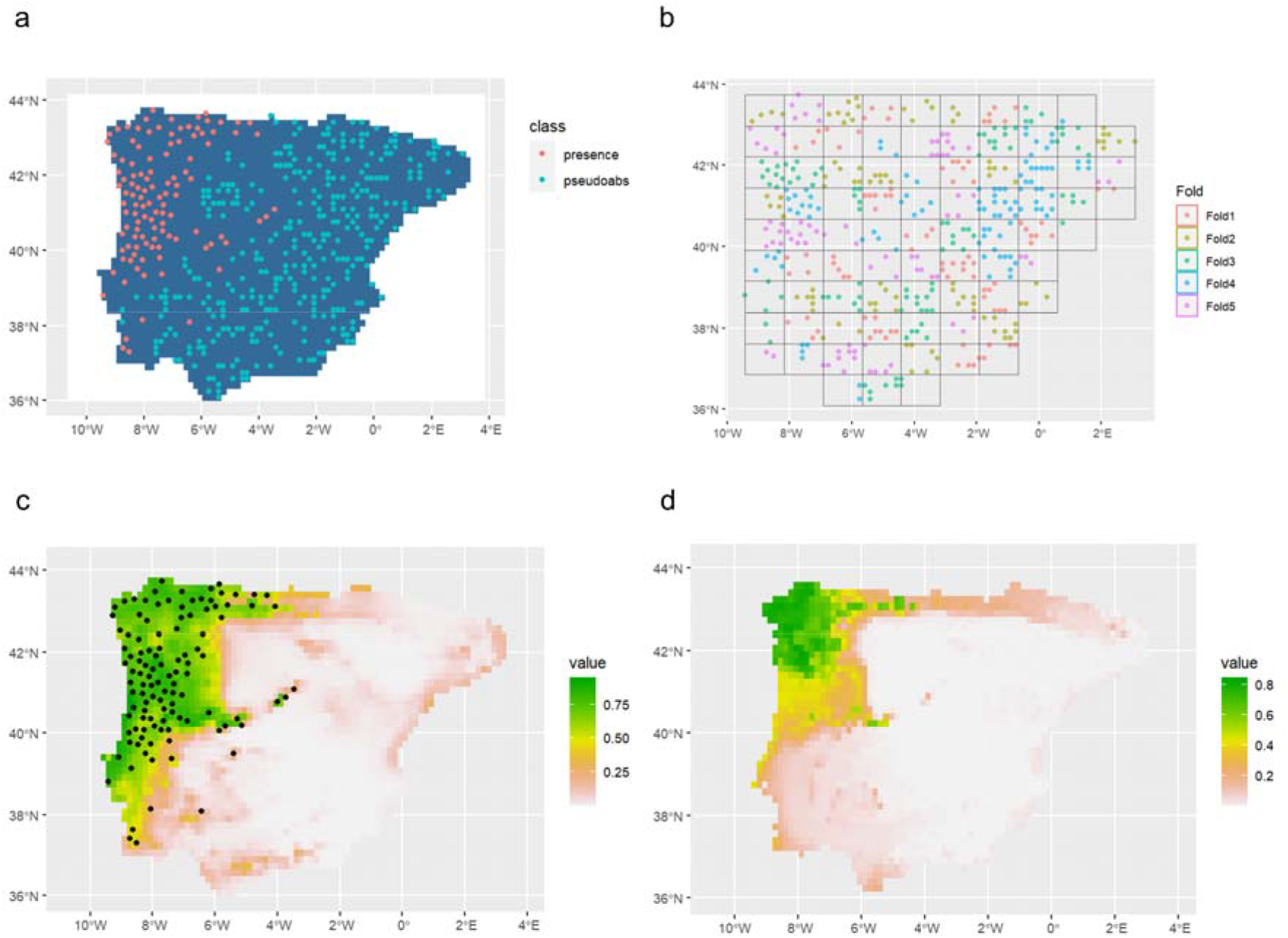
outputs of the SDM pipeline using present-day occurrences (example 1). a) thinned presences and sampled background points; b) spatial blocks for cross-validation; c) projection of the ensemble over the present; d) projection of the ensemble over the future

The choice of environmental variables is also a very important step in SDMs. It is generally desirable to focus on variables that are of biological significance for the focal species, as that increases the ability of models to make sensible predictions for the future. Furthermore, a particular issue of concern is that environmental variables are often highly correlated, and collinearity can be an issue for several types of models (*e*.*g*. glms and gams). *tidysdm* provides functions to identify informative variables, as well as remove correlated ones. Besides relying on prior knowledge, informative variables can be identified by looking for mismatches between the presences and the underlying background (pseudo-absences) (Miller *et al*. 2021a). We can do so using violin plots to contrast the values for presences and pseudo-absences, *plot_pres_vs_bg()*, and choose variables for which presences use values different from the background. *tidysdm* also offers a quantitative approach that ranks variables based on the proportional overlap of the respective density plots: *dist_pres_vs_bg()*. Finally, the environmental variables of interest can be tested for collinearity: this is a necessary step for some models where correlated variables can lead to model instability (Guisan *et al*. 2017), and it is easily solved by setting a correlation cut-off with the function *filter_high_cor()* (note that this function provides a sensible set of uncorrelated variables, but other such sets will likely exist and might be even better, therefore we recommend the users to inspect the variables with care and make decisions based on their biological understanding of the system).

Once we have selected the presences and pseudo-absences (fig. 1a), and associated them with a set of climatic variables of interest, we are ready for modelling, and this is the step at which *tidymodels* come into play. In *tidymodels*, a *workflow* is created by combining a *recipe* to process the data and a *model specification*. The model specification can have a few selected values for the hyperparameters or can prescribe their tuning. A workflow can then be fitted and tuned to the data. This is usually done through cross-validation, typically using a form of spatial blocking to generate data folds that account for the autocorrelation intrinsic to spatial data (Figure 1b). *tidymodels*, through the additional package *spatialsample* (Mahoney *et al*. 2023), provides a simple syntax and infrastructure to perform all these steps (including additional ways of creating data folds, e.g. by clustering points based on similarity in the environmental parameters). The major advantage of *tidymodels* is that, by simply changing the model specification, it is possible to fit different models with the same code defining the rest of the *workflow*. The standardisation of tuning and fitting *workflows* allows us to take advantage of a further abstraction via *workflowsets*, which provide an easy syntax to create multiple workflows that differ in their *recipe* or *model specification* but are fitted to the same data folds. This allows us to explore multiple algorithms without having to retype a lot of code, allowing for simple-to-read code with fewer chances of error when changes are made to the pipeline (as a change will apply to all workflows automatically). *tidysdm* provides both functions to simplify some of these steps (such as automatically generating model specifications with tuning for hyperparameters usually studied in SDMs) as well as expanding some methods from *tidymodels* that are not compatible with *sf* objects.

Model tuning requires metrics to assess the goodness of fit of different hyperparameter sets. The package *yardstick* (Kuhn *et al*. 2023) is used in *tidymodels* to provide various standard metrics, such as the Area Under the Curve (AUC, (Swets 1988)); *tidysdm* uses the *yardstick* infrastructure to add metrics commonly used in SDMs (e.g. the maximum True Skill Statistics (TSS, (Allouche *et al*. 2006)), the Boyce Continuous Index (BCI, (Hirzel *et al*. 2006)), and Kappa (K, (Cohen 1960)). Because of the standard grammar provided by *yardstick*, it is possible to explore multiple variables simultaneously, and *tidysdm* offers a standard set of metrics for SDMs via *sdm_metric_set()*. An important assumption made by *yardstick* is that, in classification models (as used in SDMs), the first level of the response variable is the event of interest; in SDMs, we are interested in presences, so the response variable needs to be a factor with presences as the first (reference) level. The function *check_sdm_presence()* provides a simple programmatic check to make sure that the variable has been formatted correctly.

In many SDM analyses, the best-performing models (based on a metric of choice, e.g. Boyce Continuous Index) are often combined into an ensemble (Araújo and New 2007) which is often used to predict future ranges. This can be done following different procedures. The easier procedure is building the ensemble using the best-performing version of each model. In *tidysdm*, we term this a *simple_ensemble* (in *tidymodels*, there is also the option to use stacked ensembles, see Couch and Kuhn 2023). The simplest way to build an ensemble is simply to start from a *workflowset*; add_member() will then choose and extract the best hyperparameter values from the tuned *workflowset*. However, it is also possible to assemble an ensemble from individual *workflows* (*e*.*g*. for very computationally demanding models where individual workflows might be fitted separately on a computer cluster). The ensemble is then ready for making predictions: *predict_raster()* takes a raster of environmental predictors and returns the ensemble prediction (by default, the mean of the models within the ensemble, but it is also possible to have the median as well as weighted versions of the mean and median). The ensemble can also be subset to include only the best models specifying a minimum threshold in the metric (e.g. Boyce continuous index, shown in Figure 1c). Finally, *tidysdm* also offers the option of creating a *repeated_ensemble* to explore the role of stochasticity in thinning and sampling presences and pseudo-absences. The simplest route is simply to loop over the code to create a *simple_ensemble*, adding each iteration to a list, which is then converted into a *repeated_ensemble* object.

*tidysdm* also offers an easy and intuitive way to convert the predicted probability of occurrence to binary format (presences against absences): first, an optimal threshold based on a given metric is calibrated with *calib_class_thresh()*, then the prediction can be performed again with *predict_raster()* specifying *type=“class”* and the desired threshold. In the vignette, *predict_raster()* is used both to predict the present range, as well as making future projections for a given Shared Socio-economic Pathways (SSP) (Figure 1d). For convenience, we rely on *pastclim* to download and manipulate the WorldClim data, but the user can bring in their own environmental data, which can be easily read with *terra*.

## EXAMPLE 2: SDM PIPELINE WITH TIME-SCATTERED OCCURRENCES

We have found that, when fitting SDMs with many existing R packages, the interfaces are often designed to automatically associate points with a single raster. However, this may be problematic in many instances, especially for palaeontological/archaeological data for which it is desirable to use time series of palaeoclimatic reconstructions. *tidysdm* is designed to handle this task with ease. We illustrate this option by modelling the niche of wild horses (*Equus ferus*) dated between 22,000 and 8,000 years BP (Figure 2a, extracted from Leonardi *et al*. 2022). This vignette is also accessible through the website at the link https://evolecolgroup.github.io/tidysdm/articles/a1_palaeodata_application.html.

**Figure 2:**
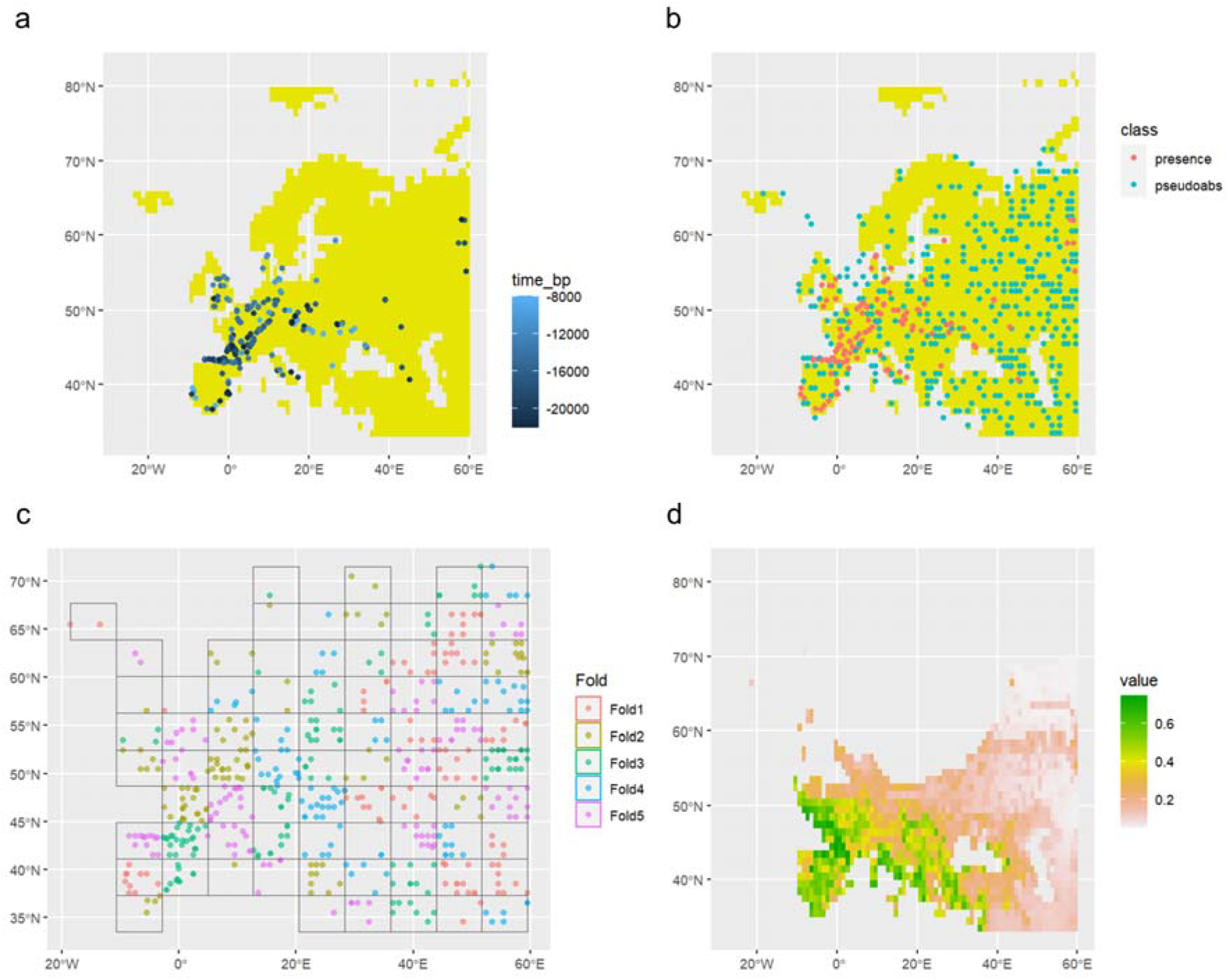
outputs of the SDM pipeline using time_scattered occurrences (example 2). a) Presences included in the analyses; b) thinned presences and sampled background points; c) Spatial blocks for cross-validation; d) Projection of the ensemble over the Last Glacial Maximum (LGM, 21,000 years BP)

For computational speed, we use a coarse time series (the Example dataset in *pastclim*), with reconstructions every 5k years and a spatial resolution of 1 arc-degree; note that much better reconstructions can be downloaded with *pastclim*. Many steps are equivalent to the ones that we described in the first example; in the next few paragraphs, we focus on the functions that are specific to time series. For this task, *tidysdm* integrates well with *pastclim*, but, in principle, the user could bring in their own *terra* SpatRasterDataset in which each variable is a *terra* SpatRaster with a time variable. *pastclim* simply makes it easier to download and manipulate the data.

The thinning of presences can be achieved with functions that are equivalent to the ones that we saw earlier, but we now need to consider also the time element. *thin_by_dist_time()* allows us to set the minimum distance in space (*dist_min*) and in time (*interval_min*) between presences. There is then the possibility to thin by keeping only one presence per cell and time slice with *thin_by_cell_time()*. For sampling pseudo-absences, *sample_pseudoabs_time()* allows the sampling of points in a manner that matches the availability of presences through time (i.e. matching sampling effort on the time axis, with the various spatial strategies available for *sample_pseudoabs()*, figure 2b). Associating presences and pseudo-absences to the correct time steps of the SpatRasterDataset can be tricky, but *pastclim* provides functions to do that.

Once we have an *sf* object with environmental variables, the process of fitting models is identical to the one described for contemporary data (example 1), see for example the spatial cross-validation shown in figure 2c. It is worth noting that, for the majority of applications, we will want to remove the time columns from the *sf* object before fitting models, since time is not a direct predictor of occurrence (but it was needed to extract the climate from the correct time step).

Once we have a *simple_ensemble()*, predicting for any given time step is straightforward, as the input climate for the time step is represented as a *terra SpatRaster* (the same as we had for present and future time steps). In this example, we illustrate this step by predicting the range at the Last Glacial Maximum (LGM, 21,000 years BP, figure 2d). *region_slice()* in *pastclim* makes it particularly easy to extract climate for any time step of interest. .

## CONCLUSIONS

*tidysdm* proves a flexible framework that leverages the power of the *tidymodels* universe, with applications for both modern species distribution/habitat suitability modelling (e.g. in ecology, conservation, biogeography, zoology) and paleo communities (including palaeoecology, archaeology and archaeozoology, palaeontology, evolutionary ecology, macroevolution, and population genetics/genomics,). A major advantage is that any addition to the *tidymodels* universe (e.g. stacked ensembles) is immediately usable for SDMs, allowing developments from the broader community of machine-learning practitioners to be quickly adopted.

## PACKAGE INSTALLATION AND AVAILABILITY

This package and the associated vignette and examples are free and open-source. It is available for download and installation on the GitHub page of the Evolutionary Ecology Group https://github.com/EvolEcolGroup/tidysdm, containing a manual, the two vignettes described here, and all details about the licence and the dependencies. We plan to release *tidysdm* to CRAN after some initial public testing. To cite *tidysdm* or acknowledge its use, please cite the present work.

## AUTHOR’S CONTRIBUTION

A.M. wrote the R package; M.L. defined the SDM pipelines with inputs from A.M. and M.C.; A.M. and M.C. wrote the vignettes with inputs from M.L.; M.L. designed the figures; all authors tested the package; A.M. wrote the function documentation and built the website; all authors wrote the manuscript.

## ACKNOWLEDGEMENTS

We thank Johanna Paijmans and Sophie Dennis for testing the package during its development.

